# A comparative study of rigid-body registration algorithms for the alignment of longitudinal structural MRI of the brain

**DOI:** 10.1101/2024.11.26.625250

**Authors:** Yazdan Rezaee Jouryabi, Reza Lashgari, Babak A. Ardekani, the Alzheimer’s Disease Neuroimaging Initiative

## Abstract

Longitudinal structural MRI (sMRI) may be used to characterize brain morphological changes over time. A key requirement for this approach is accurate rigid-body alignment of longitudinal sMRI. We have recently developed the automatic temporal registration algorithm (ATRA) for this purpose. ATRA is a landmark-based approach capable of registering dozens of serial sMRI simultaneously in an unbiased manner. The aim of the research presented in this paper was to evaluate the accuracy and inverse-consistency of ATRA in comparison to three commonly used sMRI alignment methods: FSL, FreeSurfer, and ANTS. In the absence of a ground truth, it is only possible to quantitatively determine the degree of discrepancy between two algorithms. We propose that if the discrepancy exceeds a certain threshold, the relative accuracy of the two algorithms could be determined visually. We computed the discrepancy between ATRA and each of the three other methods for the alignment of 150 pairs of sMRI taken roughly one year apart. We visually rated the accuracy of alignments in cases where the discrepancy was greater than .5 mm while the rater was agnostic to the registration method. In those instances, ATRA was considered more accurate than FSL in 46 out of 48 cases, more accurate than FreeSurfer in 6 out of 7 cases, and more accurate than ANTS in all 6 cases. ATRA was also the most inverse-consistent method. In addition to being capable of performing unbiased group-wise registration, ATRA is the most accurate algorithm in comparison to several commonly used rigid-body alignment methods.

## 1 Introduction

The brain macrostructure changes slowly with time in normal aging, as well as in certain neurological and psychiatric disorders, albeit at faster rates (Bachman et al. 2014; Ardekani et al. 2016; Goff et al. 2018). Longitudinal high-resolution 3D T_1_-weighted structural magnetic resonance imaging (sMRI) of the brain may be used to monitor brain structural changes over time. A faster than expected rate of morphological change in the brain may be indicative of unhealthy aging or abnormality. For example, (Goff et al., 2018) used longitudinal sMRI to show hippocampal volume loss during the initial eight weeks of antipsychotic treatment in first-episode drug-naive patients with schizophrenia. Whereas, no loss was detected in the age-matched control group over the same period. Furthermore, the rate of hippocampal atrophy was associated with the duration of untreated psychosis (DUP) in patients. In another study, (Bachman et al., 2014) used longitudinal sMRI to measure the rate of change of the *circularity* of the corpus callosum mid-sagittal cross-sectional area in cognitively normal (CN), mild-cognitive impairment (MCI), and early Alzheimer’s disease (AD) patients. Significant rates of reduction were found in all three groups. Moreover, the rates of change were significantly greater in MCI compared to CN, and in early AD compared to both CN and MCI groups. Subsequently, (Mubeen et al., 2017) showed that longitudinal changes in brain morphology measured by sMRI over a six-month period significantly enhanced the performance of a Random Forest classifier in predicting conversion from MCI to AD.

Since brain changes, particularly over relatively short periods of time (< 1 year), are subtle, precise image alignment is essential, whether longitudinal sMRI are compared visually or by computerized image analysis. Since longitudinal sMRI is within the same individual, the registration model of choice is rigid-body transformation in 3D with six degrees of freedom. There are several well-known algorithms developed for this purpose. Amongst these are the FSL (Jenkinson & Smith, 2001; Jenkinson et al., 2002), FreeSurfer (Reuter et al., 2010), and ANTS (Avants et al., 2008). Another recently develop method is the *automatic temporal registration algorithm* (ATRA) by (Ardekani, 2022), which is landmark-based with some important advantages over the other methods, e.g., the ability to register dozens of images simultaneously in an inverse-consistent and unbiased manner.

For the purpose of tracking longitudinal changes in brain structure, it is very important to compare the performance of these algorithms under identical conditions because even submillimeter errors in alignment could hinder accurate determination of subtle atrophy over time. The main objective of the present paper was to compare the accuracy and inverse-consistency of FSL, FreeSurfer, and ANTS in relation to ATRA in the context of alignment of longitudinal sMRI in CN older adults, MCI, and AD patients.

Representations of rigid-body transformations are dependent on the choice of basis vectors in 3D Euclidean space. Once that choice is made, the transformations may be described by 4×4 matrices that are elements of the *special Euclidean group SE* (3)(Duan et al., 2013). Unfortunately, all four algorithms (ATRA, FSL, FreeSurfer, ANTS) considered in this paper use different basis vectors in ℝ^3^. Therefore, their comparison necessitates deciphering change of bases between the four methods. This information is reported in detail in Appendix A. Given a unified formulation for the registration matrices of the four algorithms, we present a novel technique for the comparison of their accuracy using real sMRI data in the absence of a gold standard.

In addition to being a *group*, the *SE* (3) space is a *smooth manifold* of dimension six, that is, a *Lie group*. By defining a Riemannian metric on the *tangent bundle* of the manifold, it becomes possible to describe geodesic curves on *SE* (3), which provide a convenient way for averaging and interpolation between rigid-body transformations. This mathematical framework is briefly summarized in Appendix B. These operations can be utilized to impose unbiasedness, inverse-consistency, and adaptation of methods that are primarily designed to register a *pair* of sMRI (FSL, FreeSurfer, ANTS) to *group-wise* registration, that is, the simultaneous alignment of more than two sMRI into a common space.

## 2 Materials and Methods

### 2.1 sMRI data

A total of 300 3D T_1_-weighted sMRI scans from 150 participants were utilized in this study. Each participant had *two* sMRI approximately one year apart. The cohort consisted of 50 CN, 50 MCI and 50 AD patients. Subject demographics are summarized in Table 1. All images were downloaded from the Alzheimer’s Disease Neuroimaging Initiative (ADNI) database (Jack et al., 2008; Petersen et al., 2010). The ADNI was launched in 2003 as a public-private partnership, led by Principal Investigator Michael W. Weiner, MD. The primary goal of ADNI has been to test whether *serial* MRI, positron emission tomography (PET), other biological markers, and clinical and neuropsychological assessment can be combined to measure the progression of MCI and early AD.

**Table 1:** Subject demographics.

|  | **CN** (n=50) | **MCI** (n=50) | **AD** (n=50) |
| --- | --- | --- | --- |
| **Age (SD)** years | 76.6 (6.2) | 75.6 (8.0) | 75.4 (7.8) |
| **Sex (M/F)** | 26/24 | 25/25 | 25/25 |
| **ISI (SD)** days | 368 (30) | 378 (51) | 374 (74) |
| **ISI:** Inter-scan interval; **SD:** Standard deviation; **CN:** Cognitively normal; **MCI:** Mild cognitive impairment; **AD:** Alzheimer's disease. |  |  |  |

### 2.2 Discrepancy measure *d_avg_*

Assume that two different algorithms (say algorithms A and B) are used for the *same* registration task of aligning a *moving volume* (*V_m_*) to a *reference volume* (*V_r_*), yielding two *SE* (3) matrices *T_A_* and *T_B_*, respectively. If the two algorithms gave the exact same solution, then the two transformation matrices would be equal (after an appropriate change of bases) and the product of one by the inverse of the other would give the 4×4 identity matrix (*I*), that is, 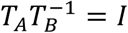. In practice, two different algorithms will inevitably yield at least slightly different results and therefore 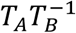 would *not* equal *I*, but 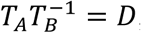, what we refer to as the *discrepancy matrix*. Since *T_A_* ∈ *SE* (3) and 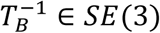, and by definition a *group* is closed under multiplication, then *D* ∈ *SE* (3), that is, the discrepancy matrix *D* itself represents a rigid-body transformation. To measure the discrepancy between algorithms A and B, we numerically apply the transformation *D* to the *brain voxels* in the moving volume *V_m_* and compute the average displacement of the brain voxels in millimeters, denoted by *d_avg_*. We will use this quantity as our measure of discrepancy between two algorithms performing the same registration task.

To compute *d_avg_* it is necessary to locate the *brain voxels* in the moving volume *V_m_*. For this purpose, we used the publicly available *acpcdetect* software from the Automatic Registration Toolbox (ART) (Ardekani et al., 1997; Ardekani & Bachman, 2009) to position *V_m_* into a standardized PIL (posterior-inferior-left) orientation. Separately, we transformed 150 skull-stripped sMRI (exclusive of the sMRI from the participants in this study) to the same standardized orientation, binarized the sMRI by setting the brain voxels equal to one and the non-brain voxels zero, and averaged them to obtain a *probabilistic brain region*. Based on this, we can infer the location of the *brain voxels* on any sMRI if it is reoriented to the standardized PIL orientation. Figure 1 shows the *probabilistic brain region* (in green colormap) superimposed on an sMRI in PIL orientation. In summary, the discrepancy measure *d_avg_* is defined as the average displacement of the *brain voxels* produced by applying the discrepancy transformation *D*.

**Figure 1:**
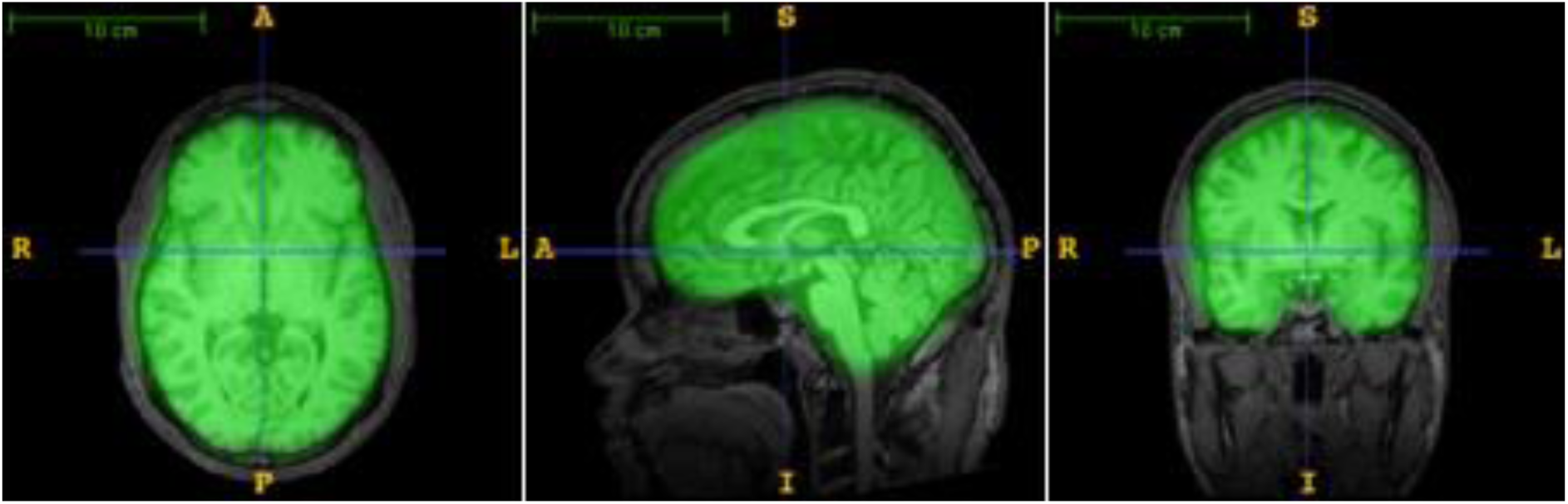
The probabilistic brain region (in green) is obtained automatically by placing an sMRI volume into a standard PIL (posterior-inferior-left) orientation. The discrepancy measure *d_avg_* is defined as the average displacement of the *brain voxels* in millimeters produced by applying the discrepancy transformation *D*.

### 2.3 Inverse-consistency

Registration of longitudinal sMRI can be considered as a process that takes a set of within-subject serial scans from different time points as input and transforms them to a new set of images with maximal anatomical alignment. In this context, the *unbiasedness* of a registration algorithm in a longitudinal setup refers to a procedure that treats *all* the images from different time points *exactly* the same. Any bias in the registration method used in a neuroimaging study will lead to a bias in all subsequent processing steps. In short, the outcome should not depend on the analyst’s “taste”.

For example, with scans at only two time points, bias is introduced by the arbitrary designation by the analyst of one scan, say the baseline scan, as the *moving volume* and the follow-up scan as the *reference volume*. Let us denote the transformation matrix thus obtained as *T* _*A*_. Now, if we reverse the role of the two scans and use the follow-up as the *moving volume* and the baseline as the *reference volume*, we obtain a second transformation matrix which we denote as *T_B_*. For an *inverse-consistent* algorithm we must have 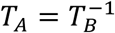, that is, the transformation obtained by reversing the roles of the moving and references volumes should be exactly the inverse of the original transformation. In practice, finite precision of machine computations means that exact inverse-consistency is not absolutely possible. In addition, some algorithms may be designed to be inherently inverse-consistent but others are not. The degree of inverse-consistency of an algorithm may be assessed by computing the discrepancy matrix, now defined as *D* = *T_B_T_A_* and the corresponding discrepancy measure *d_avg_* as the average brain voxel displacement produced by *D*.

We performed 300 registrations using each of the four registration methods (ATRA, FSL, FreeSurfer, ANTS) on our cohort of 150 participants, each participant having a baseline and a roughly one-year follow-up sMRI. For each algorithm, we first used the baseline sMRI as the *reference volume* and the one-year follow-up sMRI as the *moving volume* to perform 150 registrations. We then performed an additional 150 registrations by reversing the roles of the moving and references volumes. For each pair of registrations, we multiplied the resulting two transformations to obtain a discrepancy matrix *D* = *T_B_T* _*A*_. We computed the average brain voxel displacement *d_avg_* from each discrepancy matrix. The results are reported as a box and whisker plot for each of the four algorithms (Fig. 2).

**Figure 2:**
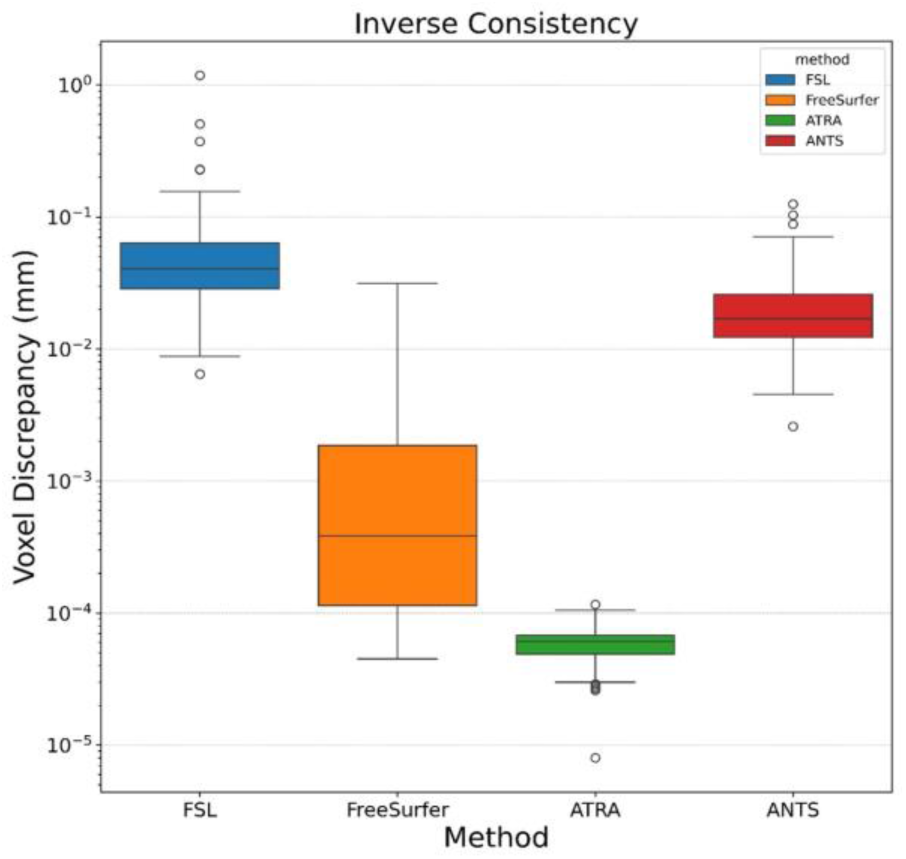
Box and whisker plot of voxel discrepancies from inverse consistency.

### 2.4 Registration accuracy

To evaluate the accuracy of registration algorithms, we randomly assigned 75 baseline sMRI and 75 one-year follow-up sMRI as the *reference* volumes and the remaining 150 sMRI as the *moving* volumes. We then registered the 150 moving volumes to their corresponding reference volumes using ATRA, FSL, FreeSurfer, and ANTS, 600 registrations in total, and obtained the corresponding *SE* (3) matrices. We then conducted change of bases in all matrices according to the procedures described in Appendix A to represent the rigid-body registration matrices of the different algorithms in a unified format and then resliced the moving volumes to match the corresponding reference volumes using the same reslicing script via trilinear interpolation. Thus, for each reference volume, we obtained 4 realigned moving volumes, one from each of the four algorithms.

To compare the accuracy of two algorithms, say algorithm A and algorithm B in a given registration task, we computed 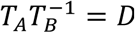 and the corresponding average brain voxel displacement *d_avg_*. If the average discrepancy was greater than .5 mm, we then visually compared pairs of realigned moving volumes (one from each algorithm). Before the visual comparison, we randomized the realigned moving volumes so that the visual rater was agnostic of the registration method. The visual rater located voxels in the pair of realigned moving volumes where there was a clear discrepancy between the results of the two methods, for example, Fig. 3a vs. 3b. This was aided by 40× magnification of the region surrounding the crosshairs displayed using pseudo colors as shown in panels 3d vs. 3e. Images were viewed in all three planes (axial, sagittal, coronal). *After* a discrepant voxel was located, the reference volume was displayed side-by-side with the crosshairs at the exact same location, colormap, and magnification (panels 3c and 3f). This enabled a decision by the rater as to which method produced a better match to the reference volume. If three such voxels were detected consequently and in all three voxels the decision was in favor and the same algorithm, then that algorithm was declared to have achieved a more accurate alignment.

**Figure 3:**
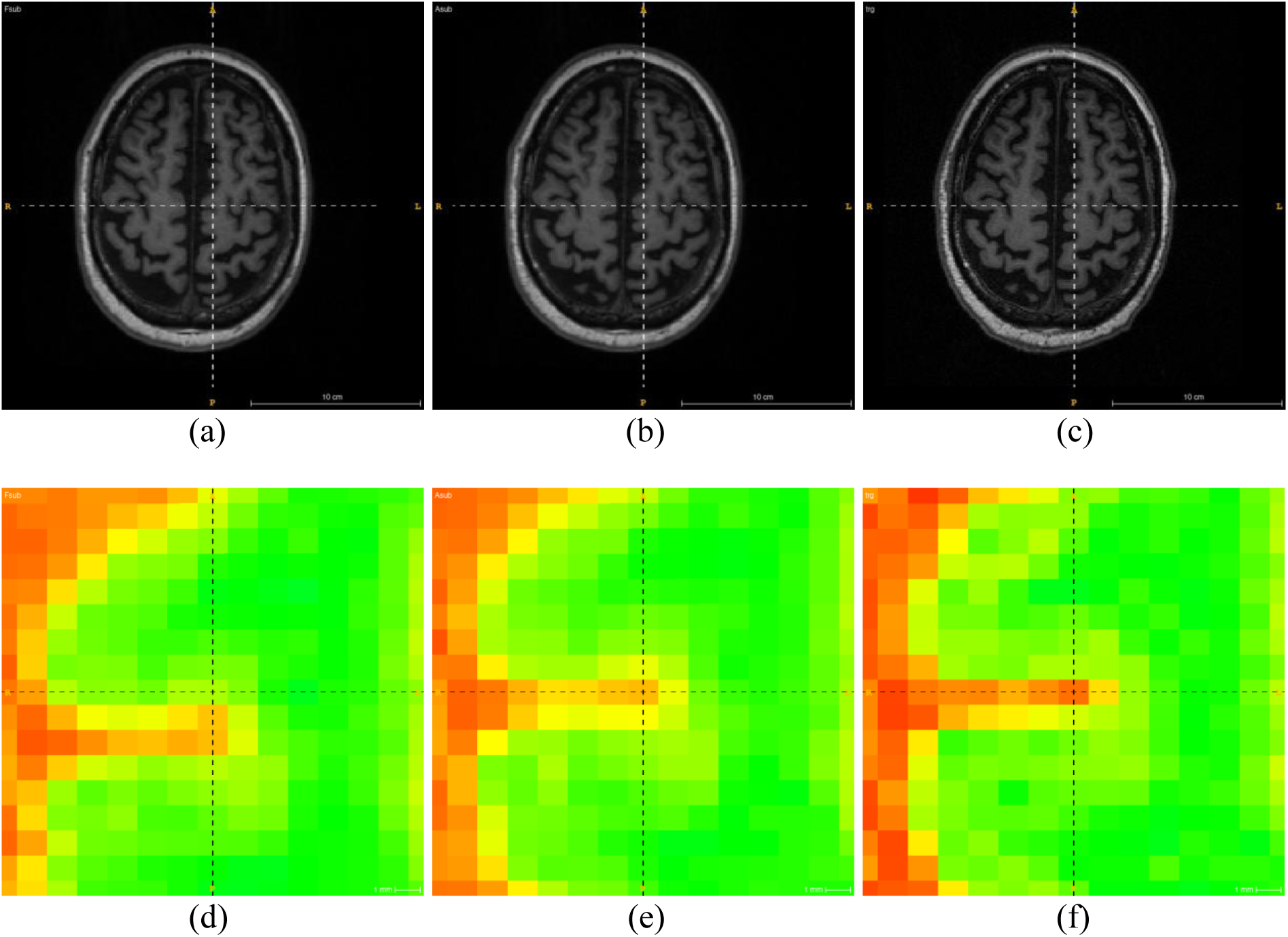
(a) and (b) Axial sections of the moving volume following alignment to the reference volume using two different methods. (c) The corresponding section of the reference volume. (d) and (e) 40× magnifications in pseudo colors of the region surrounding the crosshairs in (a) and (b). A discrepancy is noted. (f) 40× magnification of the same region on the reference slice showing a better match with (e).

In this study, we used this approach to evaluate the accuracy of the FSL, FreeSurfer, and ANTS registration algorithms in comparison to ATRA.

## 3 Results

### 3.1 Inverse-consistency

The inverse-consistency test was conducted on the four registration methods. For each of the 150 subjects, the baseline image was selected as the reference and the follow-up image as the moving volume. The resulting *SE* (3) registration matrix was denoted by *T* _*A*_. Then the follow-up volume was selected as the reference and the baseline volume selected as the moving volume. The resuting registration matrix was denoted by *T_B_*. The discrepancy matrix was computed as *D* = *T_B_T_A_* and the average brain voxel displacement *d_avg_* was computed by applying the discrepancy matrix to the brain voxels as determined by the probabilistic brain mask (Fig. 1). The 150 average displacements obtained for each of the four algorithms are shown as box and whisker plots in Fig. 2 where the vertical axis is presented on a logarithmic scale to enhance data visualization.

### 3.2 Registration accuracy

Figures 3 and 4 show two examples of the visual method for comparing the accuracy of two algorithm. Panels (a) and (b) show image slices from the moving volume resliced to match the corresponding slice of the reference volume (c) using the two registration methods. Panels (d) and (e) display 40× magnifications of a rectangular region surrounding the crosshairs in pseudo colors. The corresponding 40× magnification of the rectangular region on the reference volume is shown in panel (f). In both Figures 3 and 4, it can be clearly seen that the registration method of panel (e) more closely matches the reference volume shown in panel (f).

**Figure 4:**
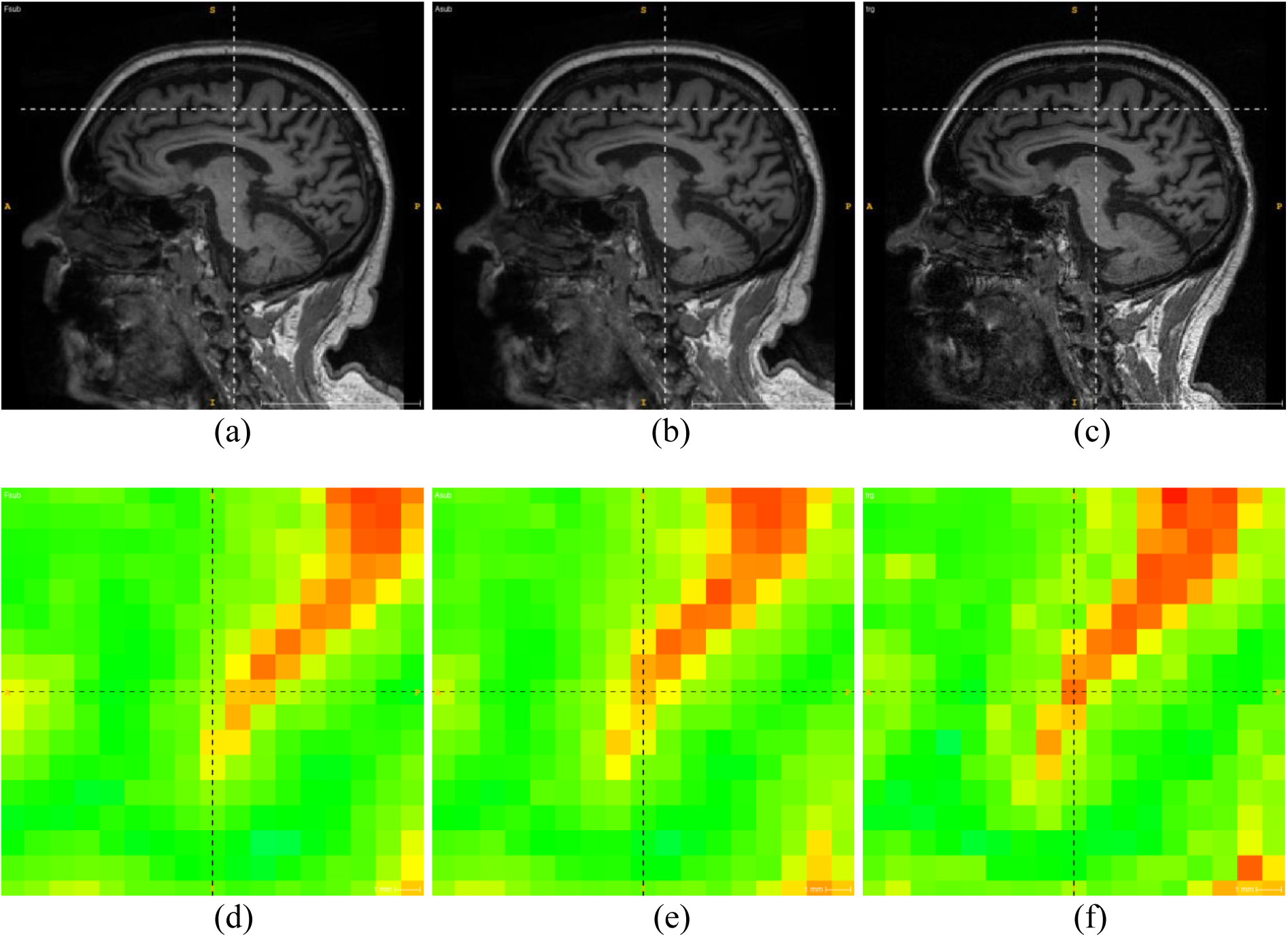
(a) and (b) Sagittal sections of the moving volume following alignment to the reference volume using two different methods. (c) The corresponding section on the reference volume. (d) and (e) 40× magnifications in pseudo colors of the region surrounding the crosshairs in (a) and (b). A discrepancy is noted. (f) 40× magnification of the same region on the reference slice showing a better match with (e).

Figure 5 shows the histogram of the 150 discrepancy measures *d_avg_* between FSL and ATRA. The visual comparison was limited to the cases where the discrepancy between FSL and ATRA was greater than .5 mm (shown in colors other than green) of which there were 48 cases (32%). After randomization of the realigned moving volumes, in 46 cases (Fig. 5, blue) ATRA was considered to be the more accurate algorithm. Only in two cases (Fig. 5, orange) FSL was considered to match the reference image more closely.

**Figure 5:**
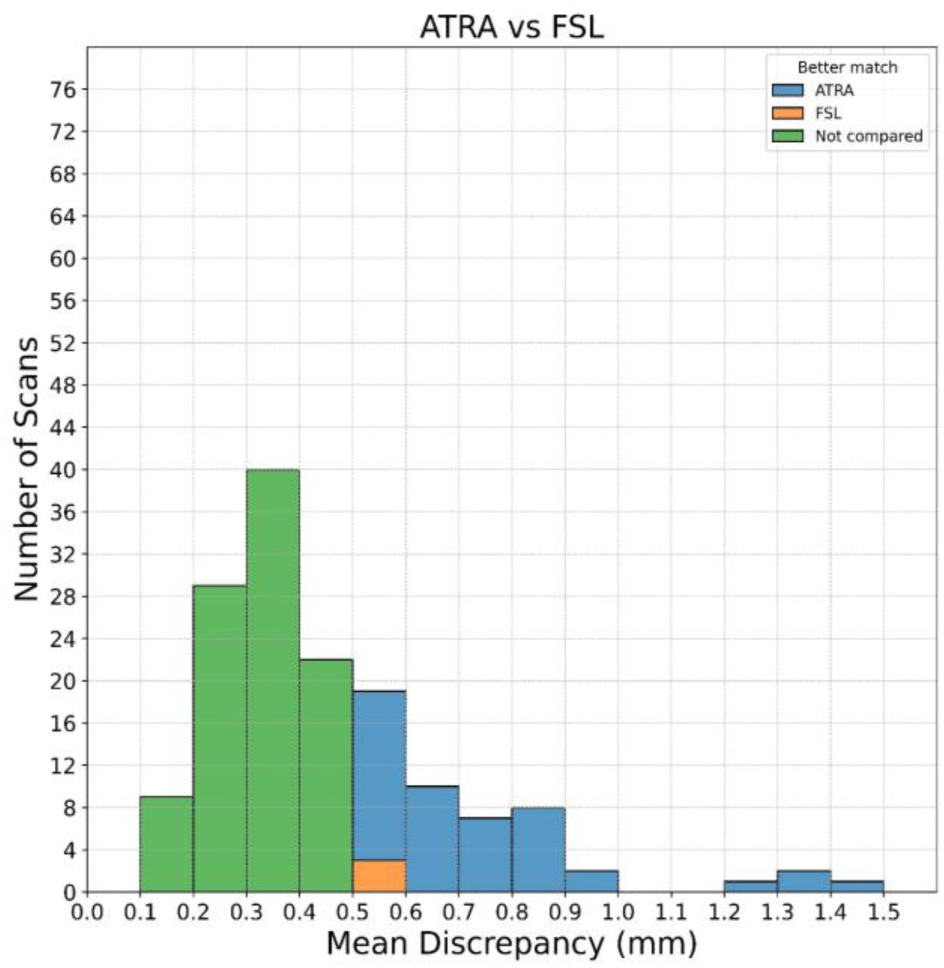
Histogram of discrepancies between ATRA and FSL registration methods. Instances where the discrepancy was less than .5 mm (102 of 150 cases) shown in green were not visually compared. Instances were ATRA was found to result in better alignment (46 cases) are shown in blue. Instances where FSL was judged a better match (2 cases) are shown in orange.

Figure 6 shows the histogram of the 150 discrepancy measures *d_avg_* between FreeSurfer and ATRA. We only visually compared the seven cases where the discrepancy was greater than .5 mm (shown in colors other than green). In six cases, the moving volume realigned using ATRA’s output *SE* (3) matrix more closely matched the reference (Fig. 6, blue). Only in one case (Fig. 6, orange) FreeSurfer was considered to match the reference more closely.

**Figure 6:**
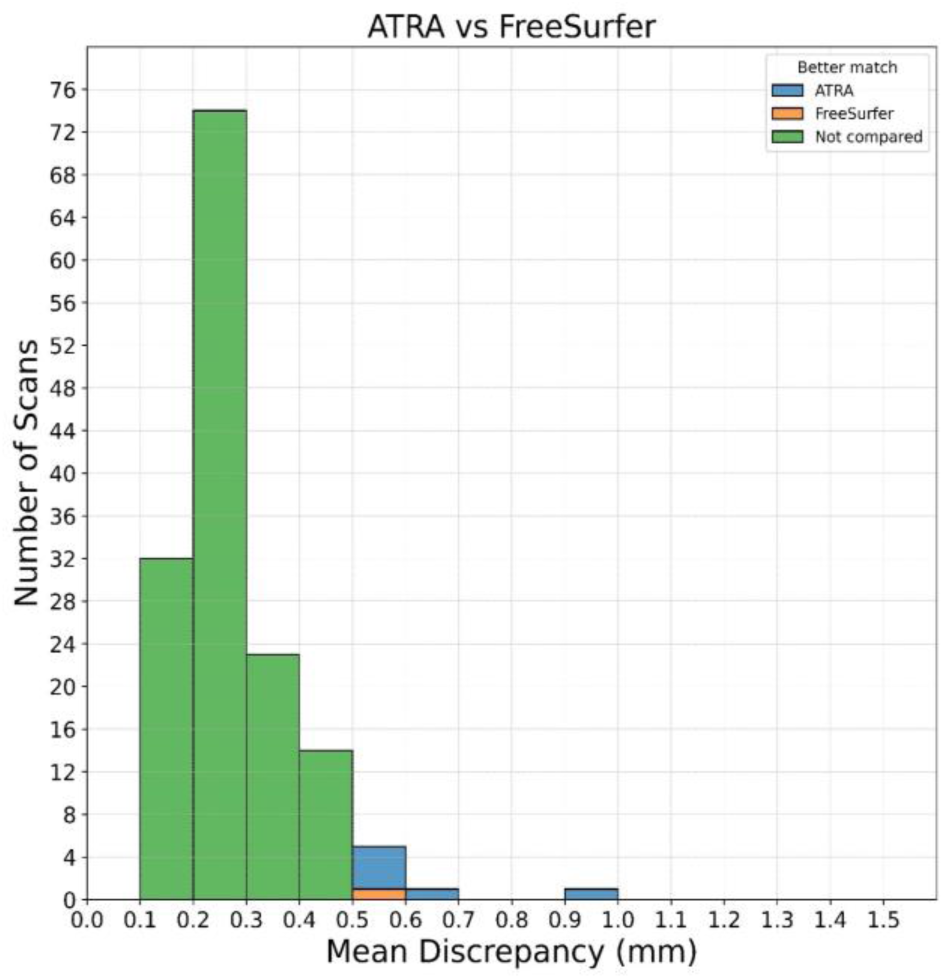
Histogram of discrepancies between ATRA and FreeSurfer registration methods. Instances where the discrepancy was less than .5 mm (143 of 150 cases) shown in green were not visually compared. Instances were ATRA was found to result in better alignment (6 cases) are shown in blue. The one case where FreeSurfer was judged a better match is shown in orange.

Figure 7 shows the histogram of the 150 discrepancy measures *d_avg_* between ANTS and ATRA. In six cases the discrepancy was greater than .5 mm. In all six cases, the moving volume realigned using ATRA’s output *SE* (3) matrix more closely matched the reference (Fig. 7, blue) as compared to ANTS.

**Figure 7:**
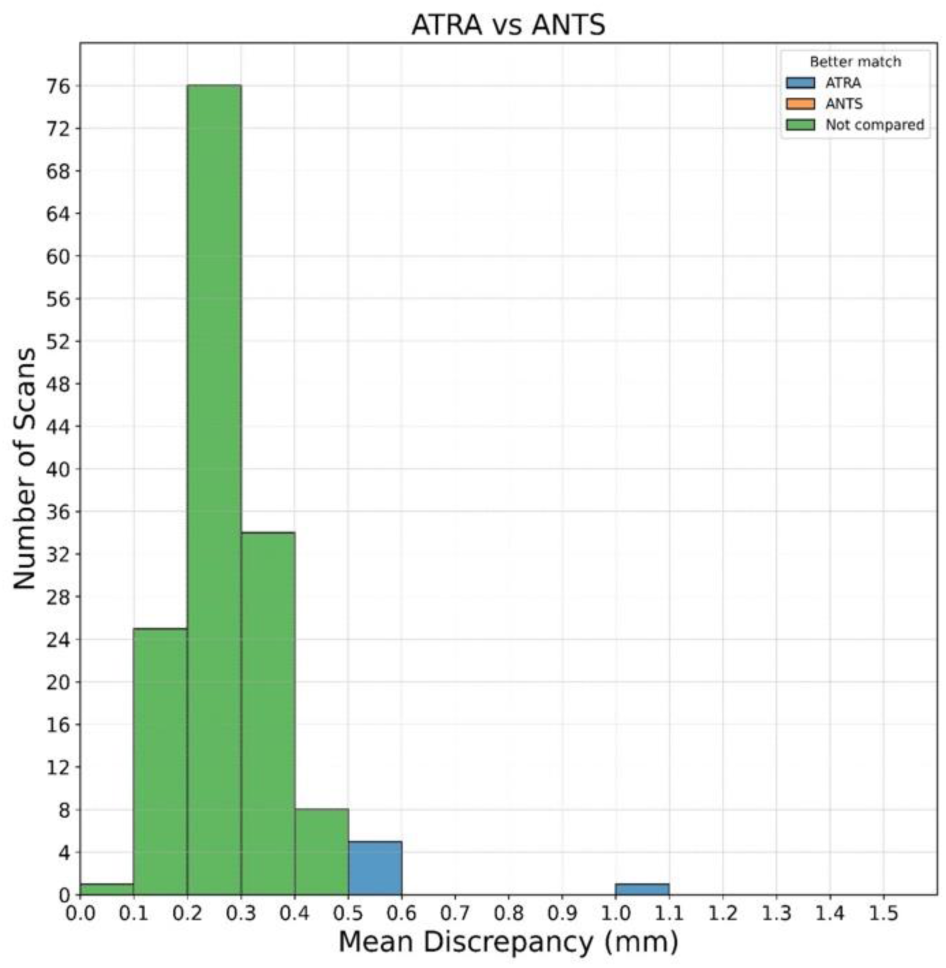
Histogram of discrepancies between ATRA and ANTS registration methods. Instances where the discrepancy was less than .5 mm (144 of 150 cases) shown in green were not visually compared. In the remaining 6 cases shown in blue ATRA was found to result in better alignment.

## 4 Discussion

Accurate spatial alignment of longitudinal intra-subject sMRI is important for applications that aim to measure subtle brain morphological changes over time. Since several public domain algorithms exist for this purpose, it becomes necessary to compare the accuracy of these algorithms. The main objective of this research was to compared the accuracy three widely used methods (FSL, FreeSurfer, and ANTS) with respect ATRA, a more recently developed landmark-based algorithm. Since the actual misalignment between longitudinal images is not known, the accuracy between different methods cannot be straightforwardly compared. However, it is easily possible to quantitatively determine the degree of discrepancy between the results of different registration methods. The working hypothesis of this research was that if the discrepancy (average voxel displacement) is greater than .5 mm, then it is possible to visually determine which algorithm more closely matches the moving volume to the reference volume.

We computed the discrepancy between FSL and ATRA for registration of 150 pairs of intra-subject sMRI acquired approximately one year apart. The histogram of discrepancies is shown in Fig. 5 ranging from .1 to 1.5 mm. In 48 cases, the discrepancy between FSL and ATRA was greater than .5 mm. After randomization of the outputs so that the rater (BA) were agnostic to the registration method, in 46 cases, ATRA was found to have produced realignments that more closely matched the reference sMRI. Interestingly, in the two cases where FSL was considered to be more accurate, the discrepancy was on the smaller side (between .5 to .6 mm). In one of the two cases, we observed significant motion artifacts in one of the images which makes both the registration and accuracy comparison more difficult.

The registration discrepancy between ATRA and FreeSurfer over the same 150 registrations is shown in Fig. 6. The results of ATRA and FreeSurfer are much closer compare to FSL. Only in 7 case the discrepancy between ATRA and FreeSurfer were greater than .5 mm. After randomization, in 6 of those cases ATRA was considered to have produced a better match to the reference volume after realignment. The one case where FreeSurfer was considered to have produced a more accurate registration was the pair with motion artifacts.

The registration discrepancy between ATRA and ANTS over the 150 registrations is shown in Fig. 7. The discrepancy between ATRA and ANTS is the smallest compared to both FSL and FreeSurfer. Only in 6 cases the discrepancy was greater than .5 mm. After randomization, in all 6 cases ATRA was considered to have produced a more accurate realignment.

Several studies (Hua et al., 2010; Westlye et al., 2009; Yushkevich et al., 2010) emphasize the importance of inverse consistency as a crucial property in preventing bias in neuroimaging research. When our aim is to register an sMRI volume 1 to an sMRI volume 2, we can take two approaches. Approach one is to designate volume 1 as the moving sMRI and register it to volume 2 taken as the reference volume. Approach two is to designate volume 1 as the reference volume and volume 2 as the moving volume and then take the inverse of the resulting *SE* (3) matrix. This essentially amounts to two different methods of using the *same* algorithm for the given registration task. Just like we computed the discrepancy between two different algorithms, one can compute the discrepancy between the two different methods of using the *same* algorithm. We computed such discrepancy for the task of registering between the pair of images for each of our 150 subjects. The results are shown as box and whisker plots for the four algorithms in Fig. 2. It can be seen that the results of ATRA are most consistent followed by FreeSurfer, ANTS, and FSL in that order. However, the inverse-consistency of all algorithms were quite good (most discrepancies being less than .1 mm). However, in a small number of cases, the inconsistency in FSL was near 1 mm which may be unacceptable.

If a registration method lacks inverse-consistency, it can be achieved by performing the registration twice switching between reference and moving volumes and “averaging” the resulting *SE* (3) matrices. However, since the space of *SE* (3) matrices is non-Euclidean, one cannot simply take the arithmetic average the two matrices. The proper method of averaging *SE* (3) matrices is to take the mid-point of the geodesic curve connecting the two matrices on a Riemannian manifold, presented in Appendix B.

When comparing longitudinal sMRI for the purpose of detecting subtle morphological changes, it is important to ensure that each sMRI undergoes one and only one interpolation operation. Often when a moving volume is registered to a reference volume, the moving volume undergoes realignment and, therefore, an interpolation, while the reference volume remains intact. This introduces a bias in registration which makes it more challenging to compare longitudinal sMRI. To ensure unbiasedness, we recommend that both the reference and moving volumes be transformed to a “halfway” point by applying the square root of the *SE* (3) solution matrix to the moving volume and the inverse square root of the *SE* (3) solution the matrix to the reference volume. The square root of an *SE* (3) matrix is the mid-point of the geodesic curve connecting the matrix to the identifity on a Riemannian manifold, presented in Appendix B.

The FSL, FreeSurfer and ANTS registration methods are designed specifically to align *a pair* of sMRI. However, currently available databases such as ADNI (Jack et al., 2008; Petersen et al., 2010), OASIS (Marcus et al., 2010; LaMontagne et al., 2019), and MIRIAD (Malone et al., 2013) comprise multiple sMRI from the same individual. In the extreme case of the Frequently Travelling Human Phantom (FTHP) dataset (www.nitrc.org/projects/fthp) 557 sMRI are collected from the same individual.

ATRA is specifically designed to simultaneously aligning dozens of sMRI, that is, to perform groupwise registration without assigning any particular sMRI as the reference. That is, the result of the groupwise alignment with ATRA will not depend on an arbitrary designation of a reference volume by the analyst. Nevertheless, pairwise registration algorithms may be adapted to perform unbiased groupwise registration using an iterative scheme whereby each volume is registered to all other volumes and the results are “averaged”. Thus, if there are *n* sMRI volumes, a given volume is selected as the reference and (*n* − 1) registrations are performed. Then the (*n* − 1) *SE* (3) matrices are “averaged” and the process is repeated for by taking each sMRI in turn separately as the reference volume for a total of *n* × (*n* − 1) registrations. The resulting *n* average matrices are then applied to realign the *n* sMRI. Then the entire process is repeated until convergence. Thus, this approach would involve *m* × *n* (*n* − 1) registrations where *m* is the number of iterations. Details of this process are presented in Appendix B.

It must be mentioned that the FreeSurfer package does provide a groupwise registration script (Reuter et al., 2012). This software, however, requires that all sMRI be of the same matrix and voxel size, which is often not the case. Therefore, if all input sMRI are not of the same matrix and voxel size, they must be made so before running the FreeSurfer groupwise registration script. However, this step itself would require an interpolation operation which would compromise image resolution. In addition, although in theory the groupwise registration algorithm of FreeSurfer is unbiased, the authors state that for computational efficiency a random volume is selected as the reference and all other volume are registered to that: “First the registration of each image … to a randomly selected image … is computed …” “Alternatively it is possible to construct all pairwise registrations and compute the average location considering all the information … This, however, significantly increases computational cost unless N [number of sMRI] is very small …”

## 5 Conclusions

This research compared the accuracy and inverse-consistency of ATRA in comparison to FSL, FreeSrufer and ANTS rigid-body registration methods for the alignment of longitudinal sMRI in normal aging, MCI and AD. Although it is not possible to evaluate the accuracy of an algorithm in the absolute sense in the absence of a gold standard, it is relatively straightforward to determine the discrepancy between the results of alignments from different algorithms. We hypothesized that when the discrepancy is greater than .5 mm, it is possible to visually compare the accuracy of alignments produced by two different algorithms when compared to the reference image. Indeed, we were able to show that ATRA produced more accurate alignments than FSL, FreeSurfer and ANTS. Furthermore, since we were able to adjudicate between algorithms with a discrepancy of .5 mm, we conclude that the registration accuracy of ATRA is at least better than .5 mm. ATRA was also found to the most inverse-consistent method, although FreeSurfer and ANTS also produced acceptable inverse-consistency. Another advantage of ATRA is that it can perform group-wise alignment simultaneously on dozens of sMRI in an unbiased way. This paper also presents methods in Appendix A for conversion between the *SE* (3) matrices produced by all four algorithms. This will be useful as reference material for future investigators and those would like to use any of the algorithms in their processing pipelines. We also discuss in Appendix B how to ensure inverse-consistency and adapt pairwise registration methods (FSL, FreeSurfer, ANTS) to group-wise registration by iteratively averaging the output *SE* (3) matrices, however, this approach increases the computational cost exponentially.

## Data statement

The sMRI data used in this manuscript were downloaded from ADNI which is publicly available. ADNI sMRI have unique image identifiers (UID). The UID of the 300 sMRI used in this manuscript are available from the corresponding author upon request. The software ATRA, FSL, FreeSurfer and ANTS are all publicly available.

## Author contributions

YRJ: Writing – original draft, Formal analysis, Software, Visualization, Investigation; RL: Writing – review and editing, Supervision, Project administration, Resources; BAA: Conceptualization, Writing – original draft, Writing – review and editing, Software, Supervision, Formal analysis, Investigation, Methodology.

## Acknowledgments

Data collection and sharing for this project was funded by the Alzheimer’s Disease Neuroimaging Initiative (ADNI) (National Institutes of Health Grant U01 AG024904) and DOD ADNI (Department of Defense award number W81XWH-12-2-0012). ADNI is funded by the National Institute on Aging, the National Institute of Biomedical Imaging and Bioengineering, and through generous contributions from the following: AbbVie, Alzheimer’s Association; Alzheimer’s Drug Discovery Foundation; Araclon Biotech; BioClinica, Inc.; Biogen; Bristol-Myers Squibb Company; CereSpir, Inc.; Cogstate; Eisai Inc.; Elan Pharmaceuticals, Inc.; Eli Lilly and Company; EuroImmun; F. Hoffmann-La Roche Ltd and its affiliated company Genentech, Inc.; Fujirebio; GE Healthcare; IXICO Ltd.; Janssen Alzheimer Immunotherapy Research & Development, LLC.; Johnson & Johnson Pharmaceutical Research & Development LLC.; Lumosity; Lundbeck; Merck & Co., Inc.; Meso Scale Diagnostics, LLC.; NeuroRx Research; Neurotrack Technologies; Novartis Pharmaceuticals Corporation; Pfizer Inc.; Piramal Imaging; Servier; Takeda Pharmaceutical Company; and Transition Therapeutics. The Canadian Institutes of Health Research is providing funds to support ADNI clinical sites in Canada. Private sector contributions are facilitated by the Foundation for the National Institutes of Health (www.fnih.org). The grantee organization is the Northern California Institute for Research and Education, and the study is coordinated by the Alzheimer’s Therapeutic Research Institute at the University of Southern California. ADNI data are disseminated by the Laboratory for Neuro Imaging at the University of Southern California.

## Data Availability Statement

The registration algorithm ATRA is available at: www.nitric.org/projects/art. Data used for the evaluation of the algorithms are available from the authors upon request.

## Appendix A

### A.1 Voxels coordinates

Each voxel in a 3D sMRI of matrix dimensions *n* _*x*_ × *n* _*y*_ × *n* _*z*_ has a set of coordinates (*i*, *j*, *k*) where *i*, *j* and *k* are integers, with *i* ranging from 0 to (*n* _*x*_ − 1), *j* ranging from 0 to (*n* _*y*_ − 1) and *k* ranging from 0 to (*n* _*z*_ − 1). The (*i*, *j*, *k*) voxel coordinates are defined by the NIFTI format and are independent of the registration method. The values of *n* _*x*_, *n* _*y*_ and *n* _*z*_ are given by the *dim* [] variable in the NIFTI file header. In this article, we use *v* to represent the (*i*, *j*, *k*) coordinates of the *moving* sMRI volume (a.k.a. *floating* or *subject* volume) written as a column vector:

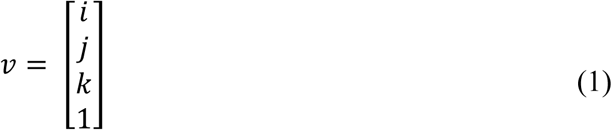

We use the prime notation to represent the voxel coordinates of the *reference* sMRI volume (a.k.a. *fixed* or *target* volume). That is:

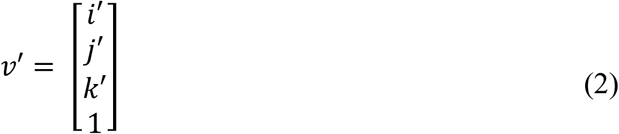

and the *reference* volume’s matrix dimensions are denoted by *n* ′_*x*_ × *n* ′_*y*_ × *n* ′_*z*_.

### A.2 Real-world coordinates

Each sMRI voxel also has the so-called real-world coordinates which we denote by (*x*, *y*, *z*) for the *moving* volume and (*x* ^′^, *y* ^′^, *z* ′) for the *reference* volume. These coordinates are usually in units of millimeters. We use *p* and *p* ′ to denote these coordinates as column vectors:

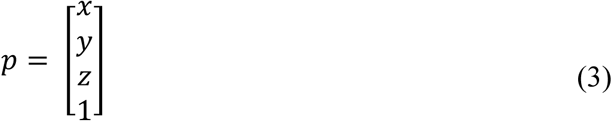

and

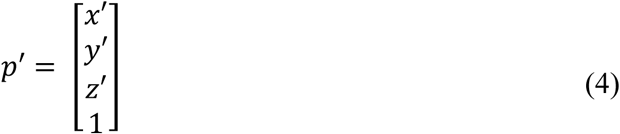

The (*i*, *j*, *k*) coordinates may be converted to the (*x*, *y*, *z*) coordinates by multiplication by an invertible 4 × 4 matrix *Q*, that is, *p* = *Qv*. Similarly, the (*i* ′, *j* ′, *k* ′) coordinates may be converted to the (*x* ′, *y* ′, *z* ′) coordinates by multiplication by an invertible 4 × 4 matrix *S*, that is, *p* ^′^ = *Sv* ′.

It is important to note that the matrices *Q* are *S* are defined *differently* by different algorithms. So, when necessary, we use subscripts to represent these matrices. For example, the matrices defined by algorithm “A” are denoted by *Q_A_* and *S* _*A*_, and the real-world coordinates are denoted *p_A_* = *Q* _*A*_*v*, and *p* ^′^ = *S* _*A*_*v* ′.

### A.3 Conversion between output transformations

Assume that we have two registration methods A and B. Their corresponding output transformation matrices may be denoted by *T_A_* and *T_B_*. These matrices map the real-world coordinates of the *moving* volume to the real-world coordinates of the *reference* volume for each algorithm, that is:

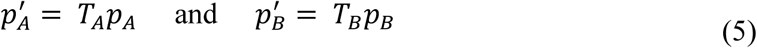

By substituting *p* ^′^ = *S* _*A*_*v* ′, *p* _*A*_ = *Q* _*A*_*v*, *p* ′*_B_* = *S_B_v* ′, and *p_B_* = *Q_B_v* in (5) we obtain:

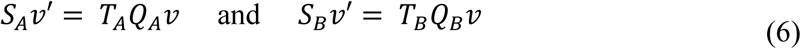

which may also be written as:

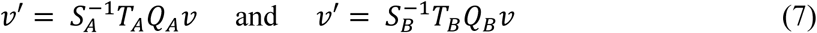

From (7) we conclude that the two algorithms A and B are equivalent when:

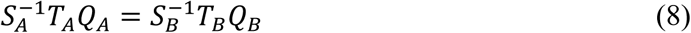

or

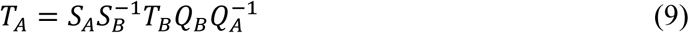

Equation (9) is the main result of this section and shows how the output transformation matrix *T_B_* from algorithm B can be transformed into a matrix *T_A_* in the format defined by algorithm A. It remains to show how different algorithms define *Q* and *S*.

### A.4 Q and S matrices as defined by ATRA

Let *d* _*x*_, *d* _*y*_ and *d* _*z*_ represent voxel dimensions in units of millimeters for the *moving* volume. These are stored from the *pixdim* [] variable in the NIFTI file header. Given these values, ATRA defines Q as:

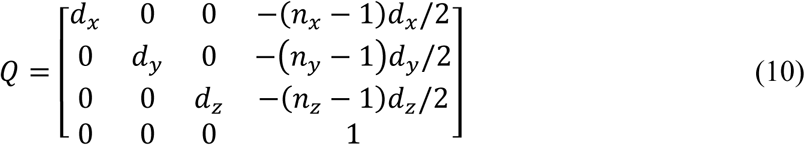

The *S* matrix is defined in exactly the same way, except we use the prime notation to emphasize that the variables relate to the *reference* volume:

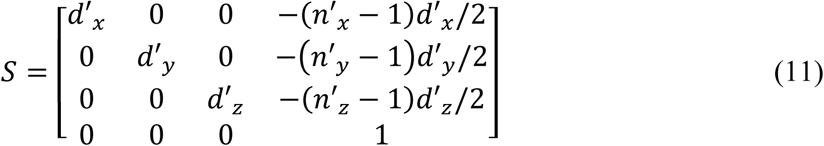

### A.5 Q and S matrices as defined by FSL

FSL actually has two slightly different methods for definition *Q* and *S*. In general, any of the axes *x*, *y* or *z* may be pointing in one of six directions with respect to the subject: right (R), left (L), superior (S), inferior (I), anterior (A) and posterior (P). When we say that the orientation of a volume is, for example RAS, it means that the *x* -axis points towards right (R), the *y* -axis points anteriorly (A) and the *z* -axis points superiorly (S). PIL would be an *x* posterior, *y* inferior, and *z* left orientation. There are 48 possible orientation codes. Half of these codes refer to a *right-handed* coordinates system (RAS, LPS, etc.). The other 24 codes refer to a *left-handed* coordinates system (PIL, RSA, etc.). For an sMRI volume in the *right-handed* system, FSL defines *Q* and *S* as follows:

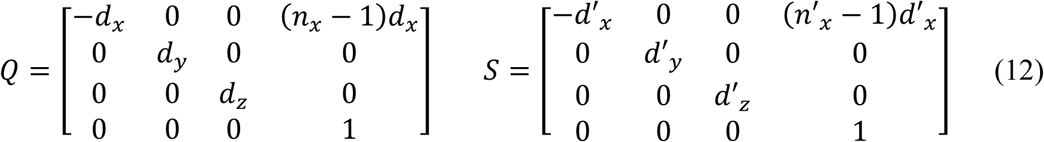

For an sMRI volume in the *left-handed* system, FSL defines Q and S as follows:

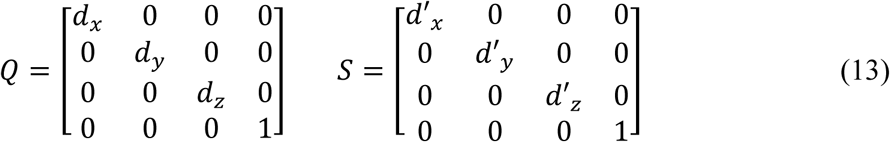

Given the definitions of *Q* and *S* in ATRA given by (10) and (11), and the definitions of these matrices in FSL given by (12) or (13), we can use the (9) to convert the transformation matrices between FSL and ATRA formats.

### A.6 Q and S matrices as defined by ANTS

The *Q* and *S* matrices used by ANTS are read directly from the NIFTI file header of the *moving* and *reference* sMRI volumes. The values are read from the *srow* _*x*, *srow* _*y*, and *srow* _*z* variables of the NIFTI file header. Given these arrays for, the *moving* volume:

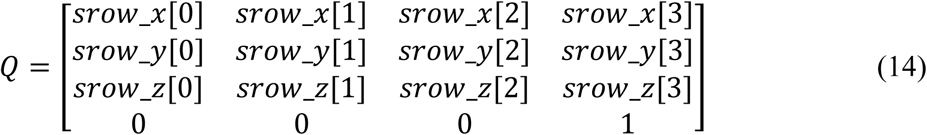

*S* is given similarly, the only difference being that the *srow* _*x*, *srow* _*y*, and *srow* _*z* arrays are read from the NIFTI header of the *reference* volume.

As we saw in (7), for a given method, we can write:

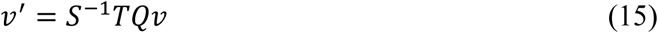

For ANTS, the *Q* and *S* matrices were defined above. The output of the ANTS method, however, does not give the transformation *T* directly in the way it is defined in (16). Some processing is needed to obtain *T* is the form that we require. Below, we will give the recipe for obtaining *T* from the output transformation of the ANTS registration algorithm.

The first step is to convert the output of ANTS into the Insight Transform File V1.0 format. This can be done using the *ConvertTransformFile* script provided by the ANTS package. The Insight Transform File contains two arrays in text format: The *Parameters* array of size 12 and the *FixedParameters* array of size 3. To obtain the *T* matrix for ANTS, we read the first 9 elements of the *Parameters* array as *R* _11_, *R* _12_, *R* _13_, *R* _21_, *R* _22_, *R* _23_, *R* _31_, *R* _32_, *R* _33_ and form a (rotation) matrix *R* as follows:

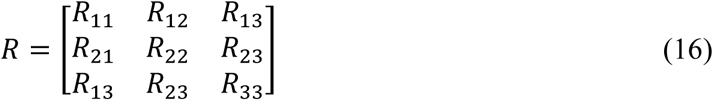

The next 3 entries of the *Parameters* array are read into a column vector *t* (translation) as:

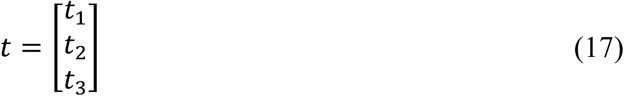

Finally, the 3 entries of the *FixedParameters* array are read into a column vector *c* (center of rotation) as:

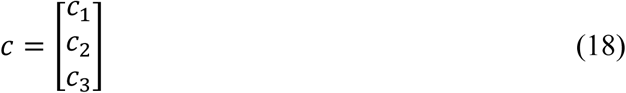

We then form a column vector *b* as:

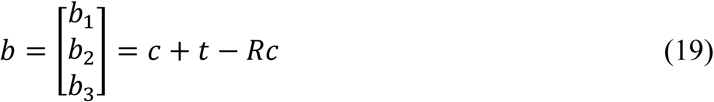

Finally, the transformation matrix *T* for ANTS may be written as:

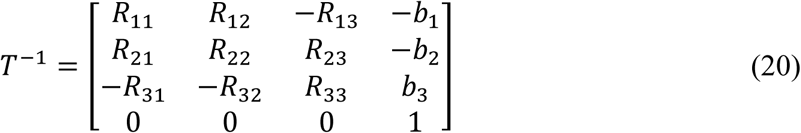

Note that in (20) the matrix *T* is given in terms of its inverse.

To summarize, we now can obtain the *Q*, *S*, and *T* matrices for ATRA, FSL, and ANTS and use our main result (9) to convert the transformation matrices between any two algorithms.

### A.7 Conversion to/from FreeSurfer

The FreeSurfer package provides the script *lta_convert* which can be used to transform the FreeSurfer output to FSL and vice versa. This allows us, therefore, to convert between FreeSurfer and any of the other 3 methods. For example, to convert the FreeSurfer output to ATRA, we first convert the matrix to FSL format and then convert from FSL to ATRA.

## Appendix B

In this appendix, we briefly cover the processes needed for Riemannian averaging on the special Euclidean group *SE* (3). These lines adopt necessities from (Duan et al., 2013), for the sake of completeness, we encourage all to refer to the main paper for a deeper understanding.

### B.1 The special orthogonal group ***SO*** (***n***)

An *n* × *n* matrix *A* is called orthogonal if 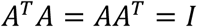. This implies that the determinant of *A* is either +1 or −1, that is, *det* (*A*) = ±1. The set of orthogonal matrices where *det* (*A*) = 1 form the *special orthogonal group SO* (*n*). These matrices represent rotations in the *n* -dimensional Euclidean space.

### B.2 The special Euclidean group ***SE*** (***n***)

Rigid-body transformation comprise of both rotations and translations. These transformations can be represented in various ways, one of which is by matrix representation. In the *n* -dimensional Euclidean space, rigid-body transformation matrices are members of the *special Euclidean group SE* (*n*) which are (*n* + 1) × (*n* + 1) matrices defined as follows:

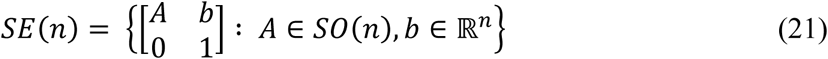

where *A* ∈ *SO* (*n*) represents a rotation, *b* ∈ ℝ^*n*^ a translation, and 0 is an *n* -dimensional row of zeroes. The outputs of all the algorithms considered in this paper are *SE* (3).

### B.3 Geodesics on the Riemannian manifold ***SE*** (***n***)

As detailed in (Duan et al., 2013), *SE* (*n*) is not only a continuous group, it is also a Riemannian manifold, that is a Lie group. Specification of a Riemannian metric on the tangent bundle of the manifold leads to the concept of a geodesic on *SE* (*n*), the shortest path on the manifold connecting two points. Specifically, let *P* ∈ *SE* (*n*) and *Q* ∈ *SE* (*n*), the geodesic from *P* to *Q* is the parametric smooth curve *γ* _*P*,*Q*_: [0,1] ⟶ *SE* (*n*) defined as follows. Let

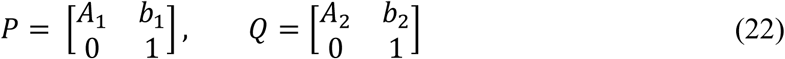

Then *γ* _*P*,*Q*_(*t*) is given by:

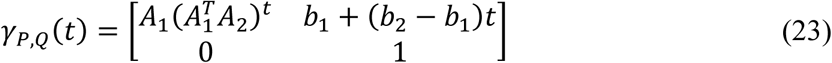

The reader can easily verify that *γ* _*P*,*Q*_(0) = *P* and *γ* _*P*,*Q*_(1) = *Q*.

### B.4 Enforcing inverse-consistency

When using a pairwise registration algorithm that is not necessarily inverse-consistent (FSL, FreeSurfer, ANTS), it is possible to enforce inverse-consistency by running the algorithm twice. Let *P* ∈ *SE* (3) be the realignment matrix obtained by registering volume A (moving) to volume B (reference); and *T* ∈ *SE* (3) be the realignment matrix obtained by registering volume B (moving) to volume A (reference). Let *Q* = *T* ^−1^. Matrices *P* and *Q* represent two different results for solving the realignment of volume A (moving) to volume B (reference). To obtain an inverse-consistent solution for this problem, we propose to simply “average” *P* and *Q*. Since these matrices reside on a Riemannian manifold, their average may be taken as the halfway point on the geodesic from *P* to *Q*. This can be obtained from (24) by evaluating *γ* _*P*,*Q*_(*t*) at *t* = 1/2, which is given by:

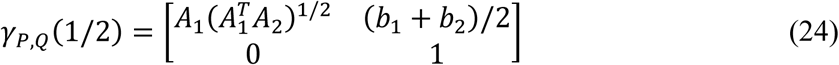

Note that the evaluation of (24) requires taking the “square root” of 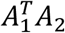. A method for performing this operation will be discussed below.

### B.5 Matrix exponential and logarithm

The *exponential* of a square matrix *A* is defined as the infinite series:

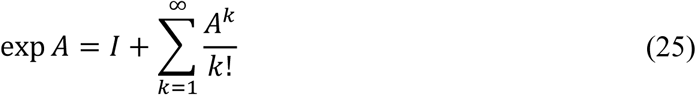

It can be shown that the series (25) converges in norm for all *A*. The inverse of exp *A* always exists and is given by:

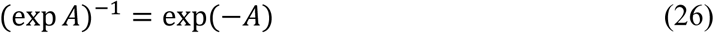

If matrices *A* and *B* commute, that is *AB* = *BA*, then:

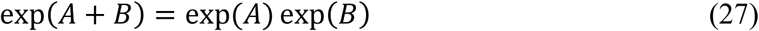

For *α* ∈ ℝ, we define:

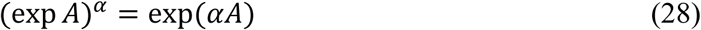

For a given matrix *B*, if there exists a matrix *A* such that *B* = exp *A*, then *A* is said to be a logarithm of *B* and we write *A* = log *B*. However, for an arbitrary *B*, the logarithm does not always exist or if it exists, it may not be unique. However, if *B* ∈ *SE* (*n*), then it can be shown that log *B* always exists and is unique. A formula for obtaining log *B* when *B* ∈ *SE* (3) is given in (Duan et al., 2013). The set of matrices:

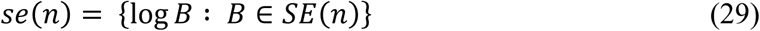

is referred to the *Lie algebra* of the special Euclidean group and is the tangent space of the *SE* (*n*) Riemannian manifold at the identity matrix *I*.

The square root of a matrix can be obtained from (28) by setting *α* = 1/2. Specifically, given a matrix *B* ∈ *SE* (*n*), we first find the unique matrix *A* = log *B* ∈ *se* (*n*) and then from (28) we have:

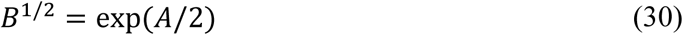

### B.6 Unbiased pairwise realignment

One of the advantages of ATRA is that the alignment brings all sMRI into a standardized common space. Every sMRI undergoes one and only one interpolation operation. The results do not depend on the order in which the images are introduced to the algorithm resulting in an unbiased realignment of all sMRI. The pairwise algorithms produce a transformation *T* ∈ *SE* (3) that aligns a moving volume to a reference volume. As such, the moving sMRI undergoes a transformation and interporation while the reference sMRI remains intact. This produces a bias in subsequent analyses. To remedy this situation, it is recommended to find *T* ^1/2^ using (30). Then transform the moving sMRI by *T* ^1/2^ and the reference sMRI by *T* ^−1/2^. Thus, both the moving and reference volumes undergo realignment to a common space.

### B.7 Groupwise realignment

An advantage of ATRA is that it produces groupwise realignment by simultaneously registering multiple sMRI. Pairwise algorithms may be adapted to perform groupwise registration at the expense of computation efficiency. Assume that there are *M* sMRI volumes to be realigned. A pairwise algorithm may be utilized to perform groupwise registration as follows:

1. Initialization step: Assume that we have *M* sMRI volumes *V* _*i*_ = 1, … *M*. Set 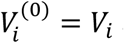 and 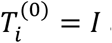 and *n* = 0
2. Repeat until convergence

2.1 Repeat for *i* = 1, … *M*

2.1.1. Perform *M* − 1 pairwise registrations to find alignment matrices *P* _*ij*_ ∈ *SE* (3) that register the moving volumes 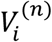 to reference volumes 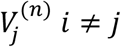.
2.1.2. Average the *M* − 1 alignment matrices *P* _*ij*_ over *j* to obtain *P* _*i*_. The algorithm for averaging *P* _*ij*_ is given in (Duan et al., 2013) and summarized below.
2.1.3. Let 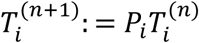
2.1.4. Apply the realignment matrix 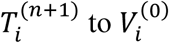 obtain 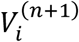
2.2. Increment *n* ∶= *n* + 1

To find the Riemannian mean of *SE* (*n*) matrices as required in step 2.1.2, let

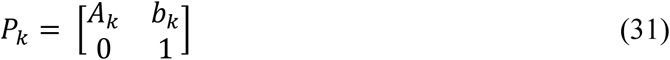

The Riemannian mean *P_avg_* of *N* matrices *P* _*k*_, *k* = 1, 2, …, *N*, on *SE* (*n*) is obtained using an iterative method based on algorithm 5 outlined in (Duan et al., 2013):

1. 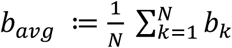
2. Set *A_avg_* = *A* _1_ as an initial input and choose a desired tolerance *ε* > 0
3. If 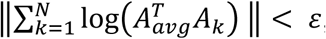, then stop
4. Otherwise update *A_avg_* as: 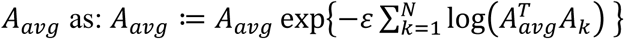 and go to step (3).

